# Exploring the infection dynamics of a bacterial pathogen on a remote oceanic island reveals annual epizootics impacting an albatross population

**DOI:** 10.1101/711283

**Authors:** Audrey Jaeger, Amandine Gamble, Erwan Lagadec, Camille Lebarbenchon, Vincent Bourret, Jérémy Tornos, Christophe Barbraud, Karin Lemberger, Karine Delord, Henri Weimerskirch, Jean-Baptiste Thiebot, Thierry Boulinier, Pablo Tortosa

## Abstract

Oceanic islands with reduced species richness provide an opportunity to investigate the emergence, maintenance and transmission of infectious diseases threatening wildlife. On Amsterdam Island, in the southern Indian Ocean, massive and recurrent mortality of the nestlings of Indian yellow-nosed albatross (*Thalassarche carteri*) has been attributed to avian cholera caused by *Pasteurella multocida*, a bacterial pathogen of likely human introduction. To understand the annual dynamics of pathogen prevalence, we measured the shedding of bacterial DNA by the albatrosses during four successive breeding seasons. The screening of 583 bird swabs by Real-Time PCR revealed an intense circulation of *P. multocida* during each study year, with a steady increase of infection prevalence across the breeding season. In the three years of highest pathogen prevalence, the epizootics were associated with massive die-offs of nestlings, inducing low annual fledging success (< 20%). These findings and developed PCR protocol have crucial applications for refining wildlife conservation plans aiming at controlling this disease.

## Introduction

Emerging infectious diseases (EIDs) are listed among the top five drivers of species extinction [1]. However, investigating infection dynamics *in natura* and identifying emergence factors are challenged by the number of animal species potentially involved in the maintenance and transmission of such pathogens. Insular ecosystems are particularly appropriate for such studies because species richness on islands, and especially on young remote oceanic islands, is generally lower than on mainland sites with comparable climate [2,3]. Furthermore, geographic isolation facilitates the identification of migratory taxa that may add complexity to a given pathosystem [4].

Amsterdam Island (37°49’S, 77°33’E) is a 53 km^2^ volcanic island lying in the southern Indian Ocean, more than 3,000 km away from any continent. In 2006, it has been designated as part of the French Southern Lands’ National Nature Reserve, to help preserving its remarkable wildlife. Amsterdam Island notably hosts several endangered seabird species, including the northern rockhopper penguin (*Eudyptes moseleyi*) and three albatross species: the Indian yellow-nosed (*Thalassarche carteri*), the sooty (*Phoebetria fusca*), and the endemic Amsterdam (*Diomedea amsterdamensis*) albatrosses [5]. Local numbers of yellow-nosed albatrosses have severely declined over the past 30 years, mirroring recurrent die-offs of the nestlings from infectious diseases [6,7], while adults do not appear to suffer from the disease [6,8]. Two microbiological studies have substantiated the link between nestling mortalities and infection by the Gram-negative bacterium *Pasteurella multocida* (*Pm*) [7,9], a bacterial pathogen known to affect wild and domestic birds worldwide [10–13]. Isolation and genotyping of the etiological agent from several hosts on Amsterdam Island supported the hypothesis of clonal infection, suggesting recent introduction of this agent [9]. The causal link between *Pm* infection and nestling mortality has been further addressed through experimental vaccination, which significantly increased nestling survival during avian cholera outbreaks [14].

The situation on Amsterdam Island thus provides a relevant framework to decipher the epidemiological dynamics of this EID, a first step towards understanding the dynamics of maintenance and transmission of a pathogen likely introduced to this ecosystem. In this study, we developed a Real-Time PCR approach allowing high throughput screening of *P. multocida* DNA to examine the infection dynamics of adult and nestling yellow-nosed albatrosses during four successive breeding seasons and in relation to the nestlings fate.

## Material and methods

### Ethical statement

The experimental design was approved by the Comité de l’Environnement Polaire (TAAF A-2013-71, A-2014-134, A-2015-107 and A-2016-80) and the French Ministry of Research (license #04939.03).

### Bird sampling

Fieldwork was conducted between December 2013 and March 2017, in the Entrecasteaux cliffs (southwestern coast of Amsterdam Island) where approximately 20 000 pairs of yellow-nosed albatrosses nest from September to March [6]. Yellow-nosed albatrosses lay a single egg in early September, that hatches between late November and mid-December [15]. Nest attendance by adults is high until January. Nestlings are then mostly on their own in their nest until fledging in April, except during feeding visits by their parents [9]. We surveyed a naturally-delineated subcolony of approximately 250 albatross pairs where bird exposure to *Pm* has been monitored since the 2013/2014 breeding season [14]. Monitored nests were georeferenced and marked with alphanumeric tags to individually identify the nestlings within a breeding season. Adults were marked with alphanumeric rings, allowing individual identification. During the four studied seasons, cloacal swabs were collected from chick-rearing adults (n=197) during the early chick rearing period using sterile applicators (one to two samples per year, per individual) and from nestlings between hatching and fledging (up to five samples per year, per individual). Swabs were conserved in 0.5 mL Longmire lysis buffer [16], kept at 0-4°C in the field, −20°C after being brought back to the base and eventually stored at −80°C until analysis. Concomitantly, nestling survival was monitored during four monthly visits to the subcolony per year, between early December (following hatching peak) and late March (before fledging). Survival monitoring was focused on the sampled nestlings, in addition to up to 30 un-manipulated (*i.e*, visually monitored without any handling) nestlings each year. Because sampling of the nestlings appeared to have no detectable effect on their survival [14], the two groups (sampled and un-manipulated) were pooled together in the survival analyses. In addition, tissue samples were collected from dead birds found on the colony. Samples were stored in 4% formaldehyde, processed and stained with haematoxylin-and-eosin for histological examination. Additional Gram staining was used for bacterial characterization (see ESM for additional details).

### Nucleic acid extraction and PCR detection of *P. multocida*

Nucleic acids were prepared from swabs using manual QiaAmp cador pathogen Mini kits (Qiagen, Courtaboeuf, France) following the manufacturer’s protocol. A Real-Time probe-based PCR protocol was developed using Pm-for (5’-ACGGCGCAACTGATTGGACG-3’) and Pm-rev (5’-GGCCATAAGAAACGTAACTCAACA-3’) primers allowing the amplification of a 116 nucleotides amplicon within KMT1 gene, a locus routinely used for the detection of *Pm* through end-point PCR [17]. Amplification was monitored in a Stratagene MX3005P (Agilent Technologies, Santa Clara, USA) thermocycler using the fluorescent Pm-probe (5’FAM-TCAGCTTATTGTTATTTGCCGGT3’BHQ1). Amplifications were performed in 25 µL final volume containing 12.5 µL of Absolute Blue real-time PCR Low Rox Mix (Thermo Scientific, Waltham, MA, USA), 0.4 µM of each primer and 0.2 µM of Pm-probe. The PCR conditions included a first Taq-Polymerase activation step (95°C for 15 min), followed by 40 cycles each composed of a denaturation (95°C for 15 sec.), an annealing (54°C for 30 sec.) and an extension (72°C for 30 sec.) step. The sensitivity of the PCR was measured by serially diluting *P. multocida* genomic DNA prepared from D2C strain [9]. The assay’s specificity was not addressed, as previous microbiological analyses have shown the occurrence of a single *Pm* clone in albatrosses from Amsterdam Island [9] and the seasonal dynamics of this clone were the focus of this study.

### Statistical analyses

Prevalence (with 95% confidence intervals) was calculated as the proportions of adults or nestlings testing positive among all sampled individuals during a given period. Variation in prevalence was quantified using logistic regressions, with *Pm*-PCR status (negative or positive) used as the response variable, and breeding season (categorical), day within the season (continuous) and their interaction as potential explicative variables. The best model was selected using Akaike Information Criterion (AIC) [18]. As many individuals were sampled several times and a large proportion of adult individuals were partners, we used generalized linear mixed models (GLMM) in the ‘*lme4*’ R package, with the individual and nest (for adults only) as random effects. Likelihood-ratio (LR) tests are reported between parentheses and odds ratios and AIC are reported in Table S2-4. Because the timing of presence in the colony differed between adults and nestlings, two distinct models were used. The proportion of nestlings surviving within the breeding seasons was quantified using the ‘*surviva*l’ package [19]. All statistical analyses were conducted in R 3.3.3.

## Results

The sensitivity of the developed RT-PCR scheme showed a positivity threshold of 9 DNA molecule templates per reaction. We screened 391 samples from 197 adults and 192 from 67 nestlings. *Pm* DNA was detected in 157/583 tested samples with some monthly prevalence exceeding 0.70. In adults, which were sampled during the early chick-rearing period only, prevalence varied significantly among breeding seasons (LR *χ*^2^ = 79, p < 0.01), reaching its maximum in 2015 (0.60 [0.50; 0.69]) and minimum in 2017 (0.01 [0.00; 0.07]; Figure 1a). In nestlings, the model with an effect of breeding season (LR *χ*^2^ = 35, p < 0.01), day (LR *χ*^2^ = 13, p < 0.01) and their interaction (LR *χ*^2^ = 7, p = 0.07) on prevalence was selected, indicating some variations among and within breeding seasons. Notably, prevalence was generally low at the beginning of the chick-rearing periods (≤ 0.14), except in 2014/2015 when prevalence was high already in December (0.70 [0.46; 0.88]). Each year, prevalence tended to increase throughout the season (see Table S1 for detailed figures). Interestingly, prevalence in nestlings was maximal in 2014/2015, 2015/2016 and 2016/2017, corresponding to seasons with very low fledging success (≤ 0.20) while it reached 0.57 [0.40; 0.81] in 2013/2014 (Figure 1b). The longitudinal survey of a subset of birds also revealed that some individuals testing positive at a given time point may test negative at one following time point (47/191 adults, and 9/50 nestlings Figure S1), suggesting that a fraction of the birds could actually clear out the infection. At the monthly sampling scale available, we did not detect an association between nestling infection status and survival (Figure S1). However, histological analyses revealed necrotic bacterial lesions in the heart, spleen and/or liver together with Gram-negative bacterial sepsis in nine of 21 necropsied albatross nestlings (Figures 2 and S2-3). This further supports the previous microbiological and experimental investigations regarding *Pm* pathogenicity [6,8,10]. Nest Pm-PCR status showed no spatial structuration at the monthly temporal scale (Figure S4).

**Figure 1.**
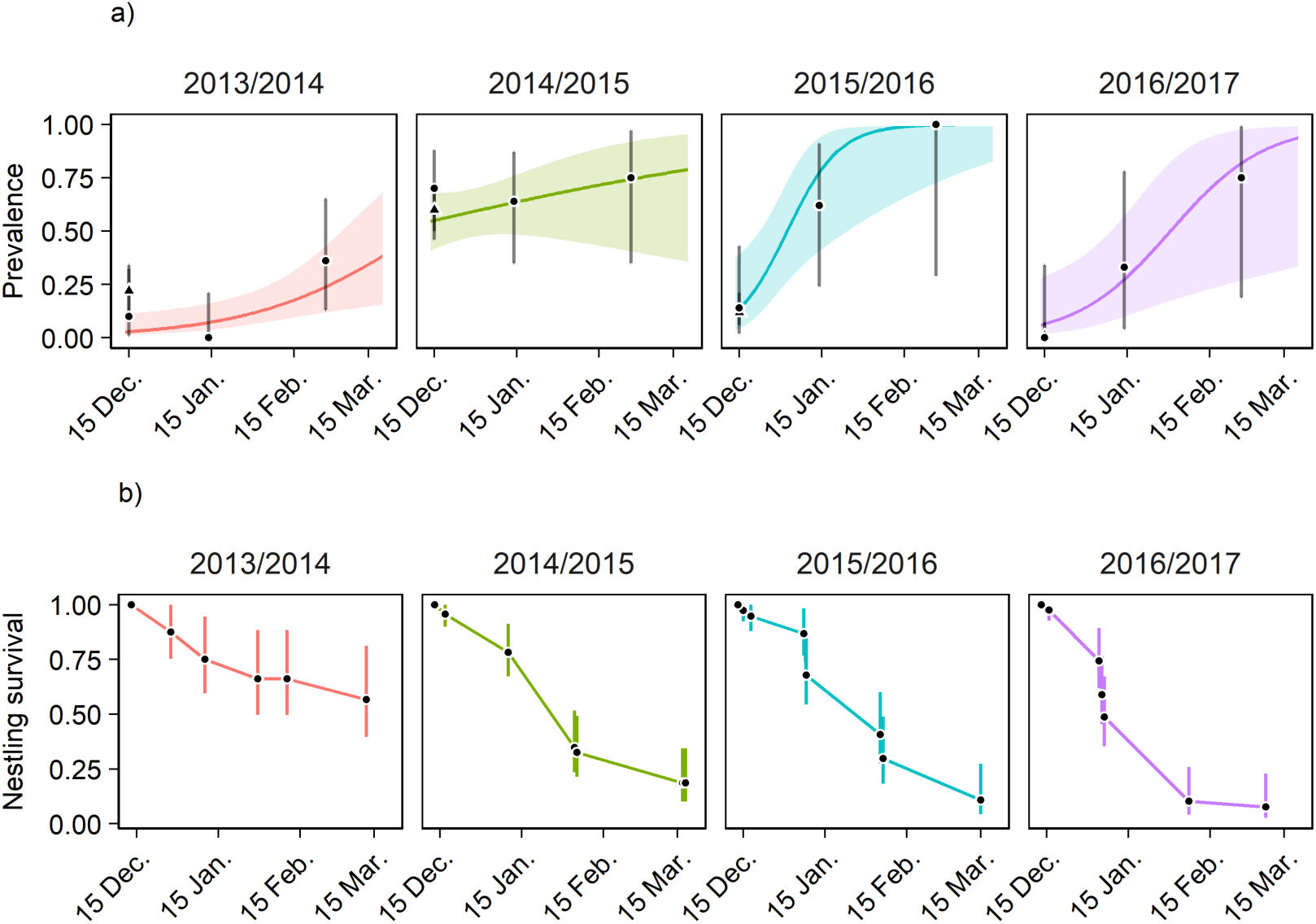
Proportion of yellow-nosed albatrosses shedding *Pm* DNA (a) and nestling survival (b) during the four breeding seasons on Amsterdam Island. Variations of prevalence over time was modelled on nestling data only; raw prevalence is represented by triangles (adults) and dots (nestlings). Bars represent the 95% Clopper-Pearson confidence intervals; shaded areas represent standard errors of model outputs; sample sizes are given in Table S1.

**Figure 2.**
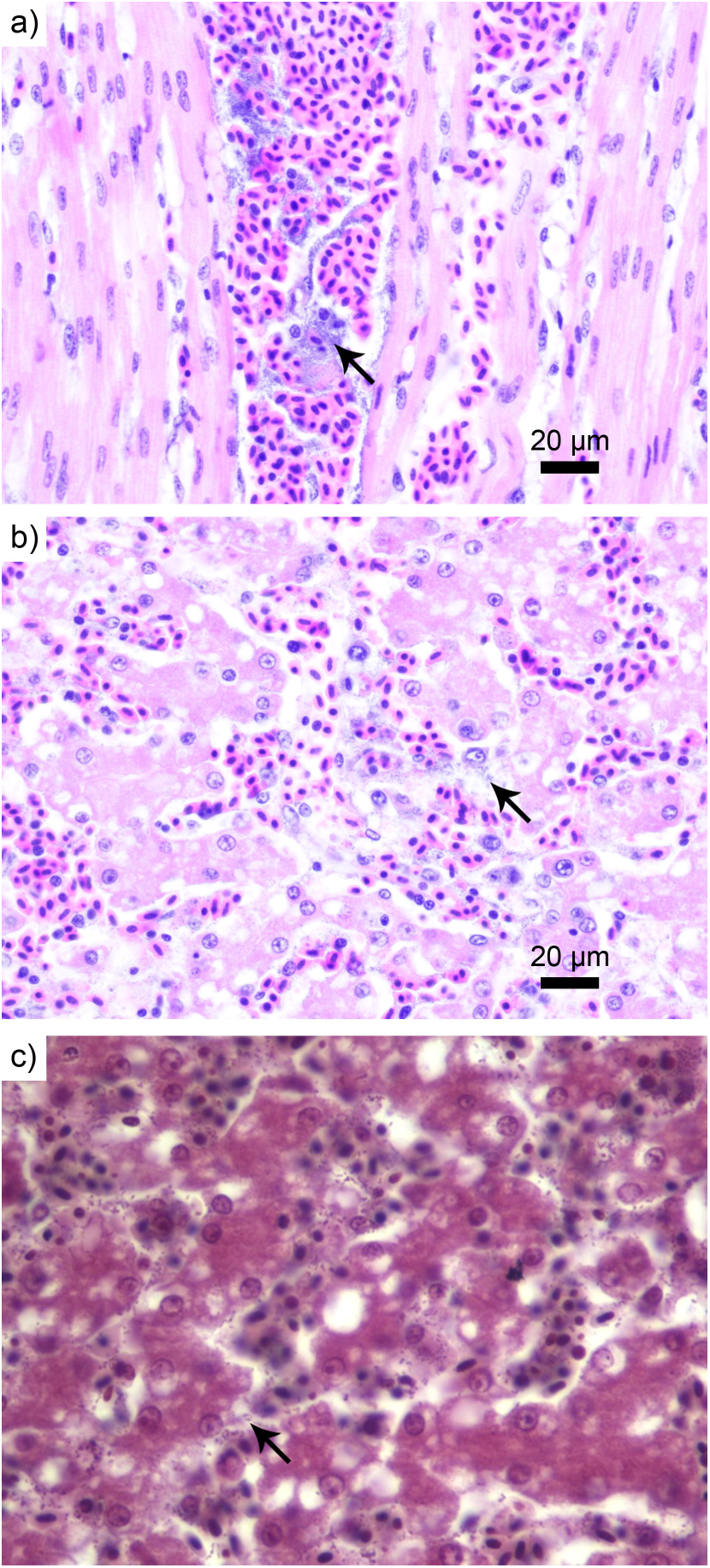
Photomicrography of a section of heart (a) and liver (b) showing circulating bacteria (black arrows). The bacteria were characterized as Gram-negative (black arrows), here in the liver (c). Haematoxylin-and-eosin staining (a and b) and Gram staining (c), magnification x 40.

## Discussion

This study reports the first multi-year investigation of *Pm* infection in an albatross population. The results reveal that yellow-nosed albatrosses on Amsterdam Island have been facing *Pm* infection at their breeding sites every year throughout the study duration. Such epizootics are in line with the massive nestling die-offs that have been recorded over the last decades on the island, especially in yellow-nosed albatrosses [7]. Infection rates were high overall, but varied among and within seasons, reaching high levels at the time when massive nestling die-offs were recorded, while adults had already left the colony. As expected considering the virulence of *Pm* [7,9,14], very few nestlings (≤ 20%) fledged in 2014/2015, 2015/2016 and 2016/2017, corresponding to the years with highest *Pm* prevalence. In contrast, in 2013/2014, prevalence was minimal, but non-null, and nestling survival exceeded 50%. Long-term monitoring would allow exploring the drivers of inter-annual variation in *Pm* prevalence, such as climatic conditions, food availability or introduced rodents’ population dynamics.

A salient pattern highlighted here is the temporality of the infection. Indeed, prevalence is most often low at the beginning of the chick-rearing period, then increases throughout the breeding season. The estimates obtained for 2014/2015 do not follow this trend, though: prevalence is already high at the first sampling point, with over 50% of the birds testing positive, and remains high throughout the season. During 2014/2015 season, the epizootic may have started earlier for reasons that remain to be elucidated. The magnitude of the infection prevalence, together with its temporal dynamics suggests that intense epizootics occur annually in this subcolony. We could not establish a direct link between infection and fledging success at the individual scale (Figure S5). This is possibly a consequence of a discrepancy between the temporal scale of the sampling (once a month) and the *almost legendary virulence* of *Pm*, which causes death within days [13] (see Figure S6 for detailed explanation). Hence, many nestlings had actually died before we had the occasion to detect them as positive. Histological analyses carried out on nestlings’ carcasses confirmed the presence of Gram-negative intravascular bacteria often associated with hepatic and pulmonary congestion, haemorrhagic myocarditis, and/or hepatic and splenic necrosis, in line with an infection by *Pm*. Considering the quickness of the epidemiological process, a finer temporal scale of sampling is needed to precisely quantify key epidemiological parameters, such as the lethality rate and the probability to clean the infection, as well as assessing the presence of chronic shedders as suggested by the longitudinal survey (see Figure S6 for detailed explanations). The results we report here will thus help to design future field studies in an iterative process [20], notably by proposing optimized designs accounting for detection probability issues [21,22].

Altogether, the results highlight the need for a comprehensive monitoring of *Pm* infection in different compartments of the host community. In particular, a monitoring scheme including the sampling of adult birds returning to the island to breed in September and the screening of introduced rats that are also potential reservoirs, would likely improve our understanding of the whole epidemiological network. Such scheme would thus help identifying the drivers of the recurrent infection outbreaks in the studied population, and evaluating the risk of spillover to the endemic Amsterdam albatross population. The novel PCR protocol we developed here is pivotal in this perspective. This research can eventually guide stakeholders to refine conservation measures and target the most relevant compartments of the pathosystem [14] for the control of severe diseases hitting endangered wildlife populations.

## Authors’ contributions

AJ, CL and PT set up the quantitative PCR. TB, CB and HW are responsible of the field research programs. AJ, TB, KD, JT, VB, JBT and AG implemented the study in the field. AJ and EL implemented the molecular analyses. KL conducted the histological analyses. AG managed the data. AG and AJ conducted the data analyses. PT and AG led the writing of the manuscript. All authors contributed to the final version of the manuscript.

## Supporting information

Electronic Supplementary Material

## Acknowledgments

We are grateful to Romain Bazire, Marine Bely, Nicolas Giraud, Hélène Le Berre, and David Hémery for their help in the field, and Cédric Marteau, Romain Garnier and Hubert Gantelet for help at various stages.

