## Supplementary material for "Exploring the infection dynamics of a bacterial pathogen on a remote oceanic island reveals annual epizootics impacting an albatross population": Electronic Supplementary Material

**Electronic Supplementary Material
S1. Supplementary text and figures**

**1. Additional methodological information**

All included birds were recruited in the frame of the present study in addition to two other specific studies: one on the effect of an autogeneous vaccine to induce a specific immune response and protection of nestlings against *Pasteurella multocida* (*Pm*; Bourret et al., 2018; Gamble et al. 2019) and one on the persistence of maternal antibodies in nestlings (Ramos et al., in preparation; follow-up of Garnier et al., 2012). Individuals included in the present study were selected using the following criteria: (1) they had not been vaccinated against *Pm* and (2) they had not received any other treatment during the duration of the study. For instance, some birds were vaccinated against Newcastle disease virus in December 2012 (*i.e.*, one year before the start of the present study) as part of the Ramos et al. (in preparation) study (following the protocol described by Garnier et al., 2012). Some other birds were parents of nestlings that were vaccinated against *Pm* (Bourret et al., 2018), but were not vaccinated themselves, and were sampled before or soon after vaccination of their nestlings. It is thus unlikely that vaccination of the nestlings could have had any impact on the probability of infection of their parents by the time of sampling (which would not be the case later, e.g., if vaccination limits shedding or increase survival without limiting shedding). These parents of vaccinated nestlings were included in the *Pm* PCR data exploration but obviously not in the nestling survival analyses. Other birds were part of the control groups (NaCl-injected or unmanipulated, *i.e.*, observed without any handling) of the *Pm* vaccination study (Bourret et al., 2018; Gamble et al., 2019).

**2. Temporal variations of *Pm* prevalence**

**Table S1.** Prevalence of *Pm* in Indian yellow-nosed albatrosses of Amsterdam Island (37°49’S, 77°33’E). *Pm­* infection was determined by PCR on cloacal swab samples (see main text). Number of individuals testing positive are indicated between parentheses; Clopper-Pearson confidence intervals between brackets. Note that February and March samples were grouped on Figure 1 a in order to increase sample size and late November samples were grouped with December samples.

| *Breeding season* | *Month* | *Stage* | *Prevalence* |
| --- | --- | --- | --- |
| 2013/2014 | December | Adults | 0.22 (15/67) [0.13; 0.34] |
| 2013/2014 | December | Nestlings | 0.10 (2/20) [0.01; 0.32] |
| 2013/2014 | January | Nestlings | 0.00 (0/16) [0.00; 0.21] |
| 2013/2014 | February | Nestlings | 0.14 (2/14) [0.02; 0.43] |
| 2013/2014 | March | Nestlings | 0.36 (4/11) [0.11; 0.69] |
| 2014/2015 | December | Adults | 0.60 (70/117) [0.50; 0.69] |
| 2014/2015 | December | Nestlings | 0.70 (14/20) [0.46; 0.88] |
| 2014/2015 | January | Nestlings | 0.64 (9/14) [0.35; 0.87] |
| 2014/2015 | February | Nestlings | 0.75 (6/8) [0.35; 0.97] |
| 2014/2015 | March | Nestlings | 0.50 (1/2) [0.01; 0.99] |
| 2015/2016 | December | Adults | 0.12 (11/91) [0.06; 0.21] |
| 2015/2016 | December | Nestlings | 0.14 (2/14) [0.02; 0.43] |
| 2015/2016 | January | Nestlings | 0.62 (5/8) [0.24; 0.91] |
| 2015/2016 | February | Nestlings | 1.00 (3/3) [0.29; 1.00] |
| 2016/2017 | December | Adults | 0.01 (1/73) [0.00; 0.07] |
| 2016/2017 | December | Nestlings | 0.00 (0/9) [0.00; 0.34] |
| 2016/2017 | January | Nestlings | 0.33 (2/6) [0.04; 0.78] |
| 2016/2017 | February | Nestlings | 1.00 (1/1) [0.03; 1.00] |
| 2016/2017 | March | Nestlings | 0.67 (2/3) [0.09; 0.99] |

**Table S2.** Selection of the models explaining the temporal variations of *Pm* prevalence in Indian yellow-nosed albatrosses of Amsterdam Island. Variation in prevalence was modelled using a logistic regression with PCR status (negative/positive) as the response variable. Individual identity was included as a random effect, in addition to the nest for adults in order to account for a potential pair effect. Breeding season was included in the model as a categorical variable and day as a continuous variable (corresponding to the interval between the actual sampling day and November 13^th^ of the considered breeding season). Π: prevalence; Df: degrees of freedom; AIC: Akaike Information Criterion; “+” denotes an additive effect, “:” denotes an interaction. The model with the lowest AIC was selected with a threshold ΔAIC of 2; when two models had a ΔAIC < 2, the more parsimonious model was selected.

| *Stage* | *Model* | *Df* | *AIC* |
| --- | --- | --- | --- |
| Adults | Π (.) | 3 | 433.4959 |
|  | **Π (breeding season)** | **6** | **334.9387** |
|  | Π (day) | 4 | 413.3875 |
|  | Π (breeding season + day) | 7 | 334.3225 |
|  | Π (breeding season : day) | 10 | 339.1930 |
| Nestlings | Π (.) | 2 | 235.8367 |
|  | Π (breeding season) | 5 | 206.1508 |
|  | Π (day) | 3 | 227.4529 |
|  | Π (breeding season + day) | 6 | 187.9314 |
|  | **Π (breeding season : day)** | **9** | **185.1702** |

**Table S3.** Outputs from the selected generalized linear mixed model ran on the *Pm* PCR data of adult yellow-nosed albatrosses of Amsterdam Island. Individual and nest identity were included as random effects in the model.

| *ADULTS* | *Estimate ± standard error* | *Odds ratio  [95% confidence interval]* |
| --- | --- | --- |
| Breeding season 2013/2014 (intercept) | -1.85 ± 0.28 | - |
| Breeding season 2014/2015 | 2.24 ± 0.34 | 9.43 [4.88; 18.22] |
| Breeding season 2015/2016 | -0.14 ± 0.43 | 0.87 [0.38; 2.01] |
| Breeding season 2016/2017 | -2.43 ± 1.04 | 0.09 [0.01; 0.68] |

**Table S4.** Outputs from the selected generalized linear model run on the *Pm* PCR data of yellow-nosed albatross nestlings of Amsterdam Island.

| *NESTLINGS* | *Estimate ± standard error* | *Odds ratio  [95% confidence interval]* |
| --- | --- | --- |
| Breeding season 2013/2014 (intercept) | -4.53 ± 1.14 | - |
| Breeding season 2014/2015 | 4.35 ± 1.27 | 78.00 [6.47; 939.99] |
| Breeding season 2015/2016 | -0.56 ± 2.16 | 0.57 [0.01; 39.40] |
| Breeding season 2016/2017 | 0.06 ± 1.91 | 1.06 [0.03; 45.17] |
| Day | 0.03 ± 0.01 | 1.03 [1.01; 1.06] |
| Breeding season 2014/2015 : Day | -0.02 ± 0.02 | 0.98 [0.95; 1.01] |
| Breeding season 2015/2016 : Day | 0.07 ± 0.04 | 1.07 [0.99; 1.16] |
| Breeding season 2016/2017 : Day | 0.02 ± 0.03 | 1.02 [0.97; 1.07] |

**3. Longitudinal monitoring of *Pm*-PCR status**
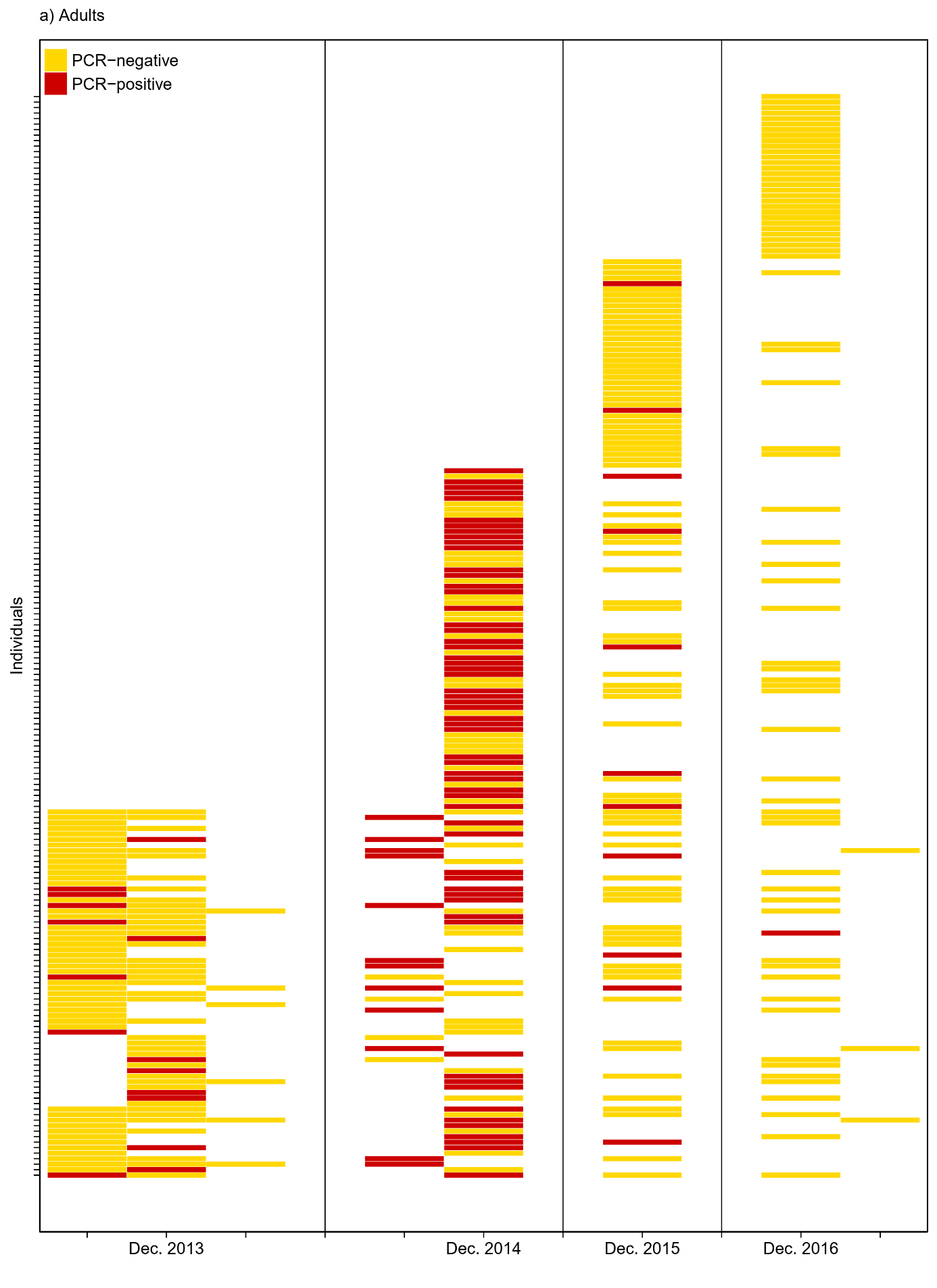

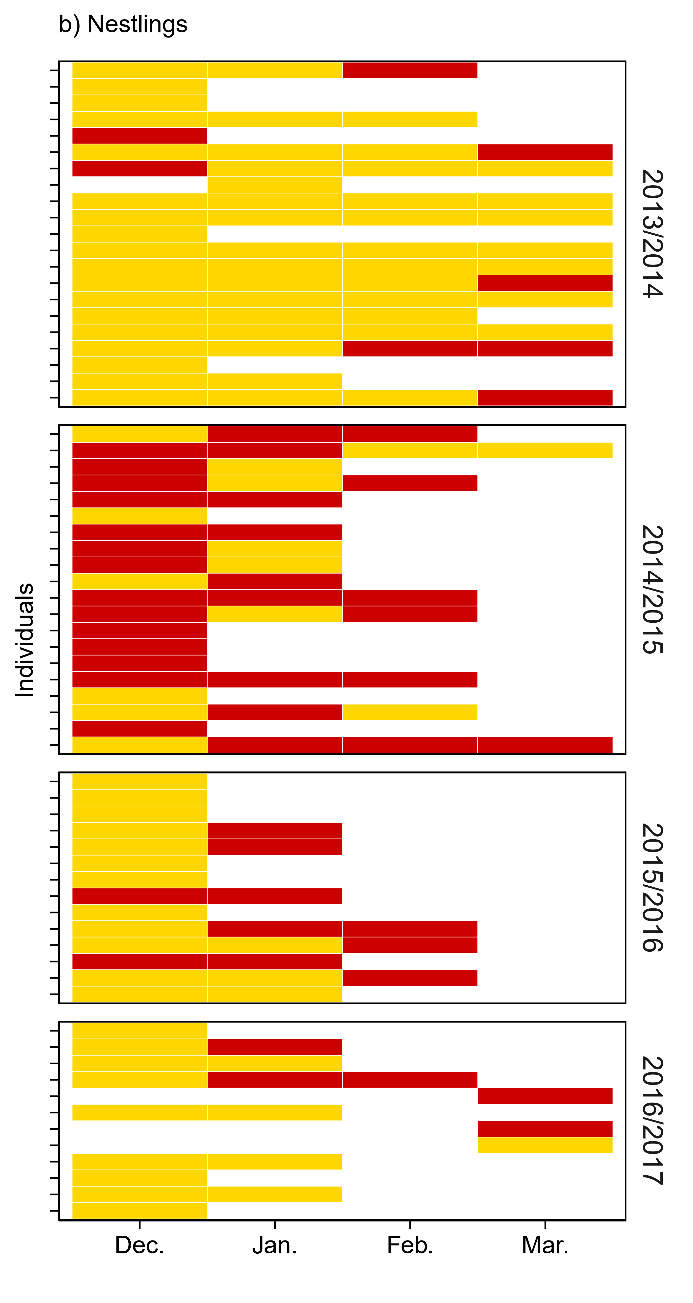

**Figure S1.** Individual histories of *Pm* detection by PCR in cloacal swab samples of adult (a) or nestling (b) Indian yellow-nosed albatrosses. Each line represents an individual and each column a month. Individuals sampled several times within a month (mostly nestlings, which were sampled twice in December) were considered PCR-positive if at least one of these samples tested positive.

**Interpretation and discussion.** Interestingly, transition from PCR-positive to -negative status was observed in 47/191 adults (191 referring to the number of individuals for which at least two samples were collected) and 9/50 nestlings (note that adults were sampled at the inter-annual scale, while nestlings were sampled at the intra-annual scale). This observation suggests that a non-negligible proportion of individuals may clean the infection. In contrast, some individuals remain PCR-positive across series of consecutive sampling occasions. It should be noted that in 2015/2016 and 2016/2017, the epizootics (mortalities of nestlings) occurred later than in 2014/2015, thus a possible rise of prevalence in adults may not have occurred without having been detected because adults were sampled before the actual outbreak. Note that adults are much less present at the colony in January and, thus more difficult to sample but also less likely to be exposed. Samuel et al. (2005) have suggested the existence of chronic carriers in wild *anatidae* populations based on antibody detection. The present study is the first to our knowledge to report molecular evidence supporting this hypothesis. If yellow-nosed albatrosses could indeed be asymptomatic carriers of *Pm*, it could have important consequences regarding spillover risks to the endemic Amsterdam albatross nesting a few kilometres away. The size of its population (less than 100 breeding pairs; BirdLife International, 2018) makes this species particularly vulnerable to potential disease outbreaks, urging the need to better understand the epidemiological dynamics of *Pm* in yellow-nosed albatrosses, the most abundant species of Amsterdam Island, but also in other species potentially involved in the maintenance and/or the dissemination of the bacterium on the island such as the brown skua (*Stercorarius antarcticus*; Boulinier et al., 2016) and the introduced brown rat (*Rattus novergicus*; Curtis, 1983). For instance, yellow-nosed albatrosses could be the maintenance host of the bacterium (Haydon et al., 2002) and brown skuas could play the role of epidemiological bridge with the Amsterdam albatross population (Caron et al., 2015). The molecular tool developed in the context of the present study will allow to efficiently investigate this hypothesis by screening other potential hosts of *Pm* on the island. Such epidemiological data, in addition to ecological data (e.g., inter-species contact rates) should help to reach the ultimate objective of describing the full transmission dynamics of *Pm* within the study system.

**4. Histology**

**Additional methodological details**

Dead birds found on the colony were necropsied within a few hours after discovery. Tissue samples from 21 yellow-nosed albatross nestlings and one adult were collected between December 2013 and January 2018. Examinations of dead birds in the study site are rare because of the high scavenging pressure imposed by brown skuas and introduced brown rats. Tissue samples were collected and stored in 4% formaldehyde at room temperature until further analysis. Samples were then routinely processed, sectionned at 4 µm thickness and stained with haematoxylin-and-eosin for histological examination. When bacteria were noticed on haematoxylin-and-eosin, additional Gram staining was performed for further bacterial characterization.

**Additional results**

Intravascular bacteria were detected in various tissues (notably liver, heart, spleen and/or lungs) of nine necropsied yellow-nosed albatross nestlings each year (see for instance Figure 2a and b, main text; Figures S2-3). These bacteria were often associated with hepatic and pulmonary congestion, haemorrhagic myocarditis, and/or hepatic and splenic necrosis. Findings are consistent with bacterial septisis as cause of death of these individuals. The bacteria were identified as Gram-negative (Figure 2c, main text), which would be in line with an infection by *Pm*.

Samples presented in Figure 2 (main text) were collected from a yellow-nosed albatross nestling found dead in the study site on the 12t^h^ January 2014 (specimen AMS13-CAD12).

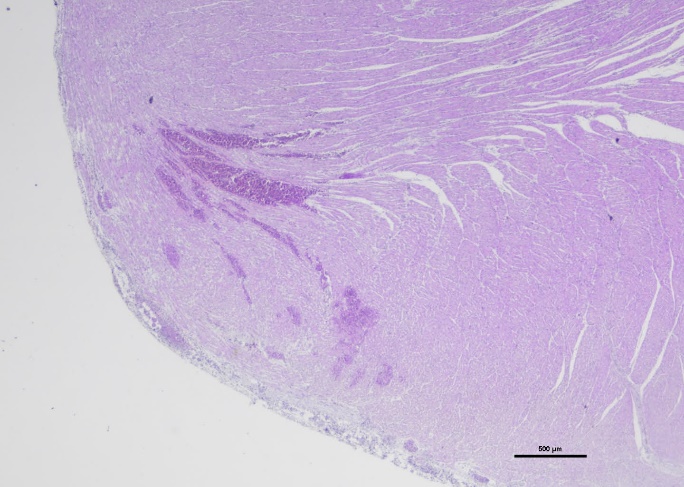

*****

a)

500 μm

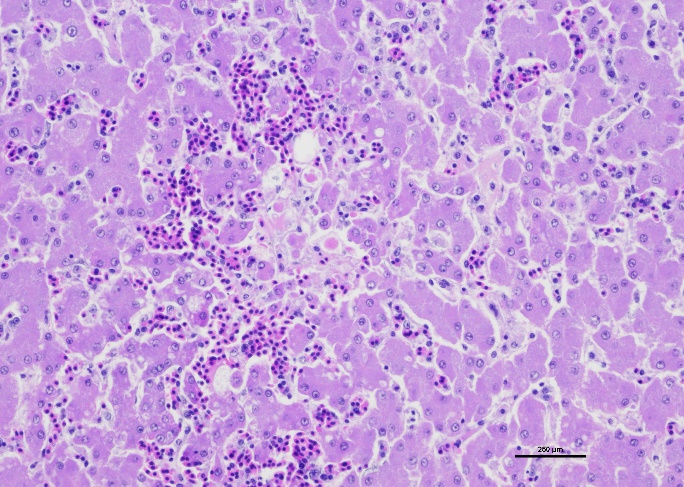

b)

200 μm

*****

**Figure S2.** Photomicrography of a section of heart (a) and liver (b) from a yellow-nosed albatross nestling (specimen AMS13-CAD11, found dead on the 11/01/2014) showing bacteria associated with haemorrhagic myocarditis and necrotic hepatitis respectively (black asterisks). Haematoxylin-and-eosin staining, magnification x 2 (a) and x 20 (b).

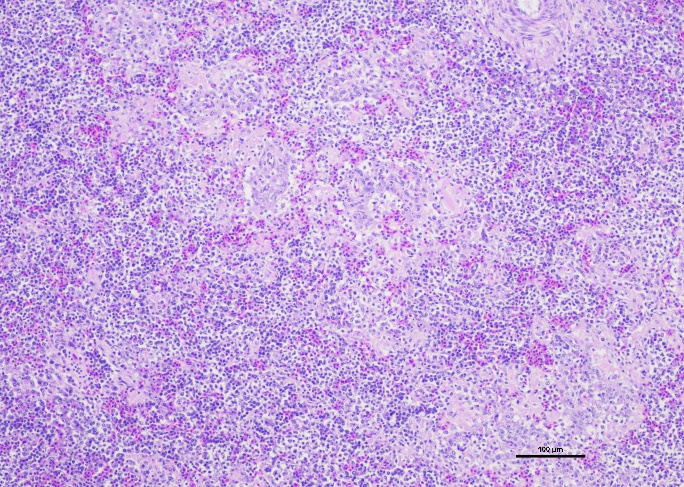

100 μm

*****

**Figure S3.** Photomicrography of a section of spleen from a yellow-nosed albatross nestling (specimen AMS16-CAD5, found dead on the 16/12/2016) showing necrotizing splenitis with intralesionnal bacteria (black asterisk). Haematoxylin-and-eosin staining, magnification x 10.

**5. Spatial patterns of *Pm*-PCR status**
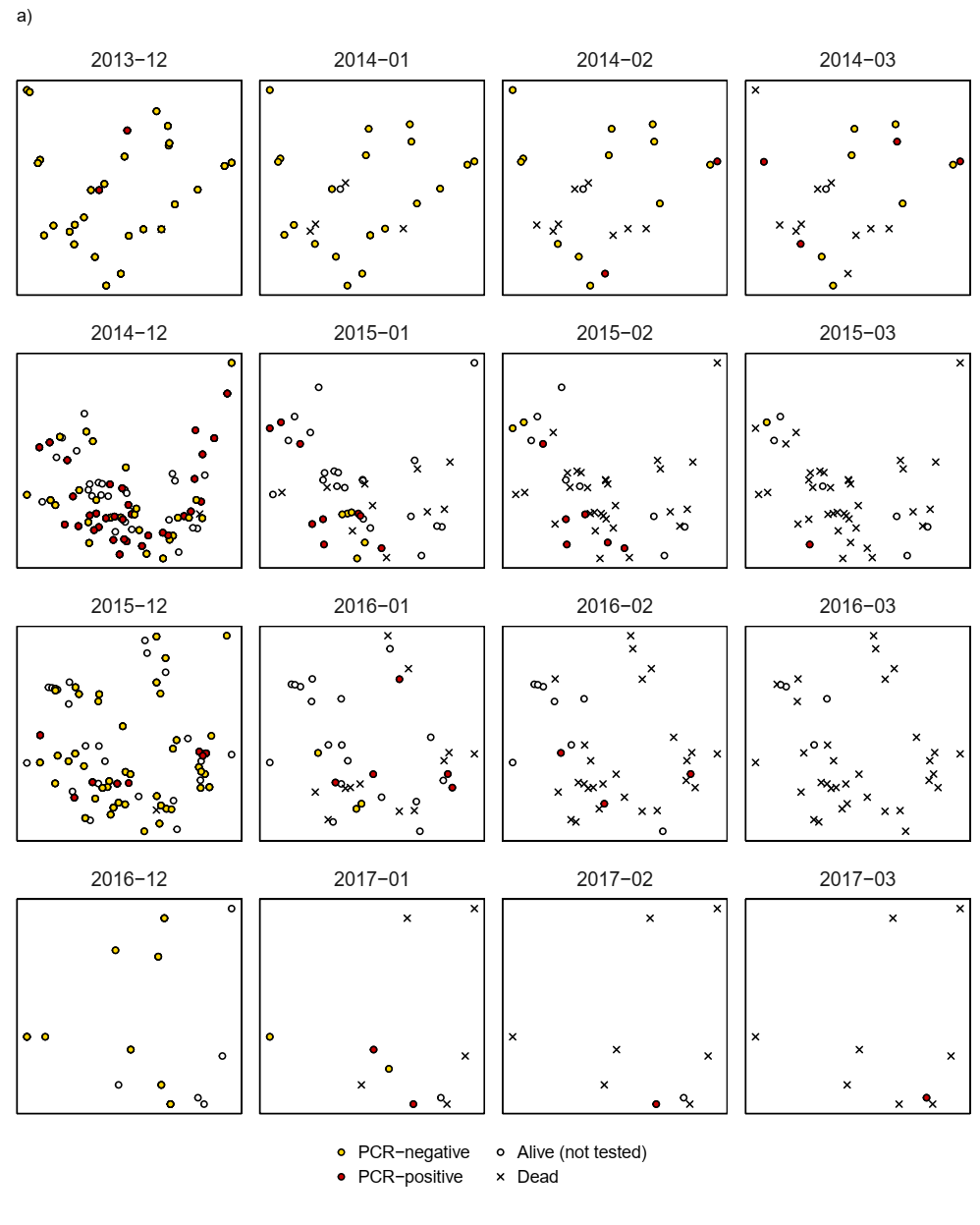

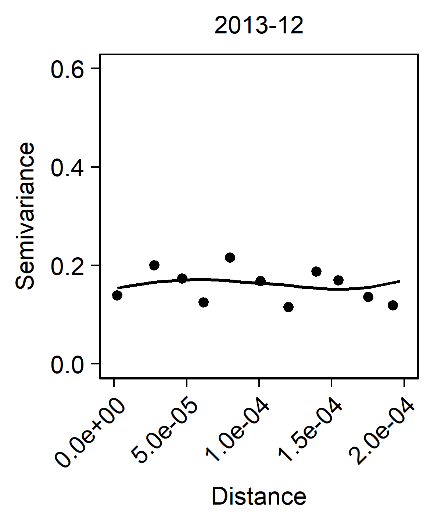

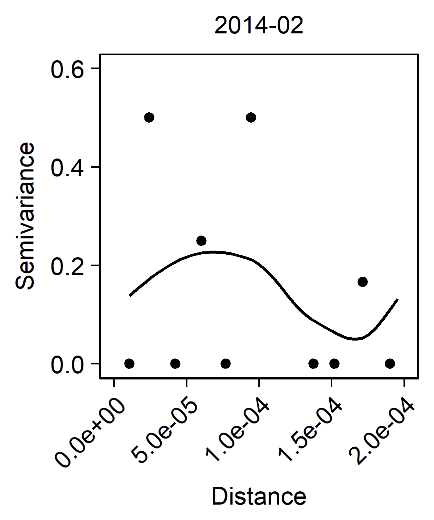

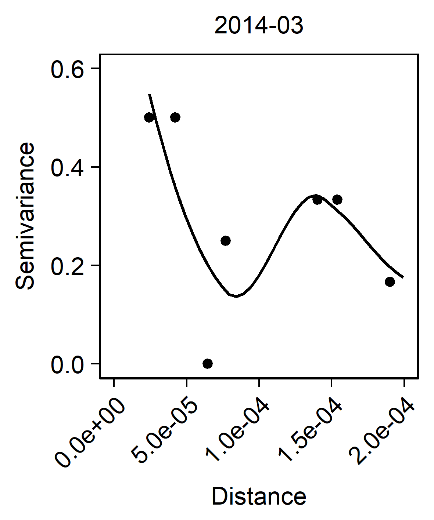

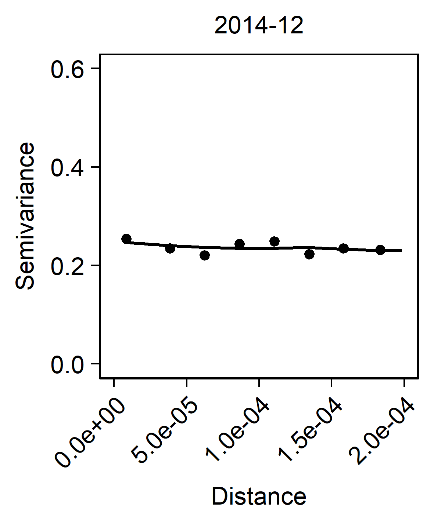

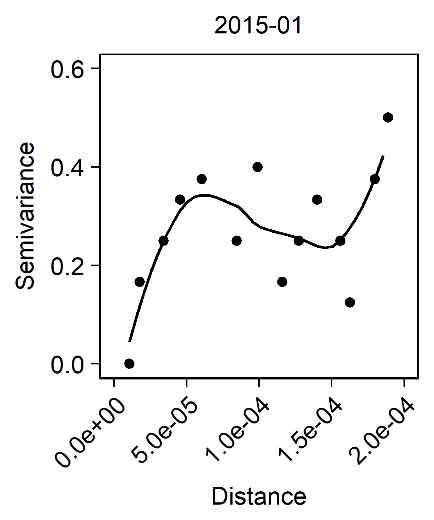

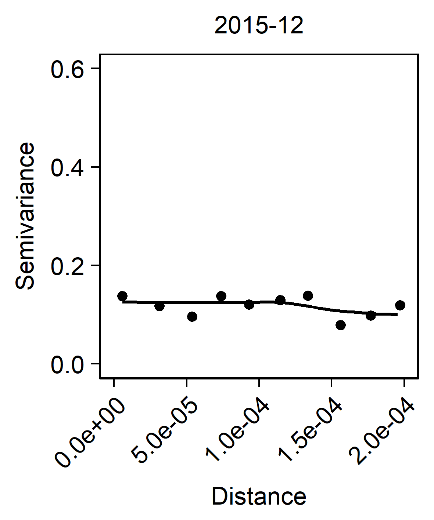

c)

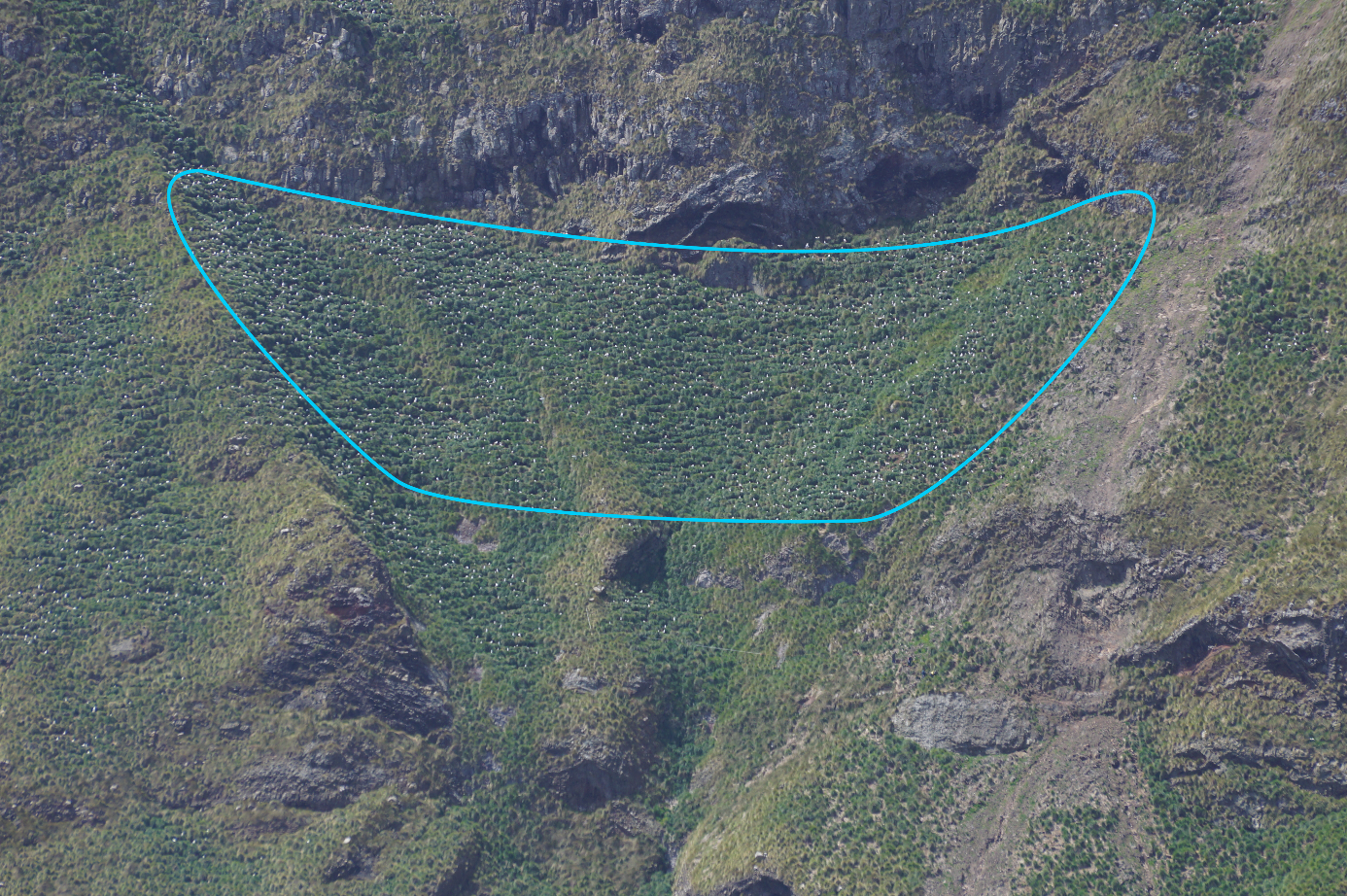

b)

See next page for figure legend (Figure S4).

**Figure S4.** (a) Spatial pattern of *Pm* detection by PCR in cloacal swab samples of Indian yellow-nosed albatrosses. Each dot represents a nest. Nests from which several individuals were tested (e.g., a parent and the nestling) within a month were considered PCR-positive if at least one of these samples tested positive. Late November samples were grouped with December samples. (b) The study colony (circled in blue) in Entrecasteaux Cliff, Amsterdam Island. Picture: Amandine Gamble/IPEV, 23 December 2016, taken from sea. (c) Variogram of the spatial distribution of *Pm* detection. Variograms were built using the in ‘*gstat*’ R package (Pebesma et al., 2004). Dots correspond to empirical variograms and solid lines to local polynomial regressions. Variograms were built only for months during which more than 10 nests were sampled and both PCR-negative and -positive results were obtained.

**Interpretation and discussion.** We could not detect any strong spatial structuration of *Pm* detection in the study colony. More precisely, the variograms of March 2014 and January 2015 are the only ones supporting the hypothesis of spatial structuration of *Pm* circulation, but in different directions (different *Pm* status between nearby nests in March 2014 *versus* similar *Pm* status between nearby nests in January 2015). Several non-exclusive transmission routes have been suggested for *Pm* such as oral and fecal shedding (Samuel et al., 2003), environmental transmission (Bredy & Botzler, 1989) or *via* rodent bites (Curtis, 1983). Characterizing the spatial patterns of *Pm* circulation could help disentangling the relative contributions of these potential transmission routes. Seabird colonies are particularly relevant models to address such questions because of their high spatial structuration at small scales. For instance, a yellow-nosed albatross nestling will not leave its nest before its first fledging attempt, four months after hatching. The spatial pattern of infection among albatross nestlings is thus a resultant of the transmission processes, and not of the sampled hosts’ movements. Few systems can allow discriminating transmission routes from sampled hosts’ movements at such a fine spatial scale. However, because *Pm* exhibits an epizootic dynamics (rapidly increasing prevalence during the breeding seasons; Figure 1a), the spatial pattern of *Pm* infection is likely to change within a breeding season. A higher sampling frequency would allow to refine the mapping of *Pm* infection.

**6. Exploration of the relationship between *Pm*-PCR status and fledging probability**

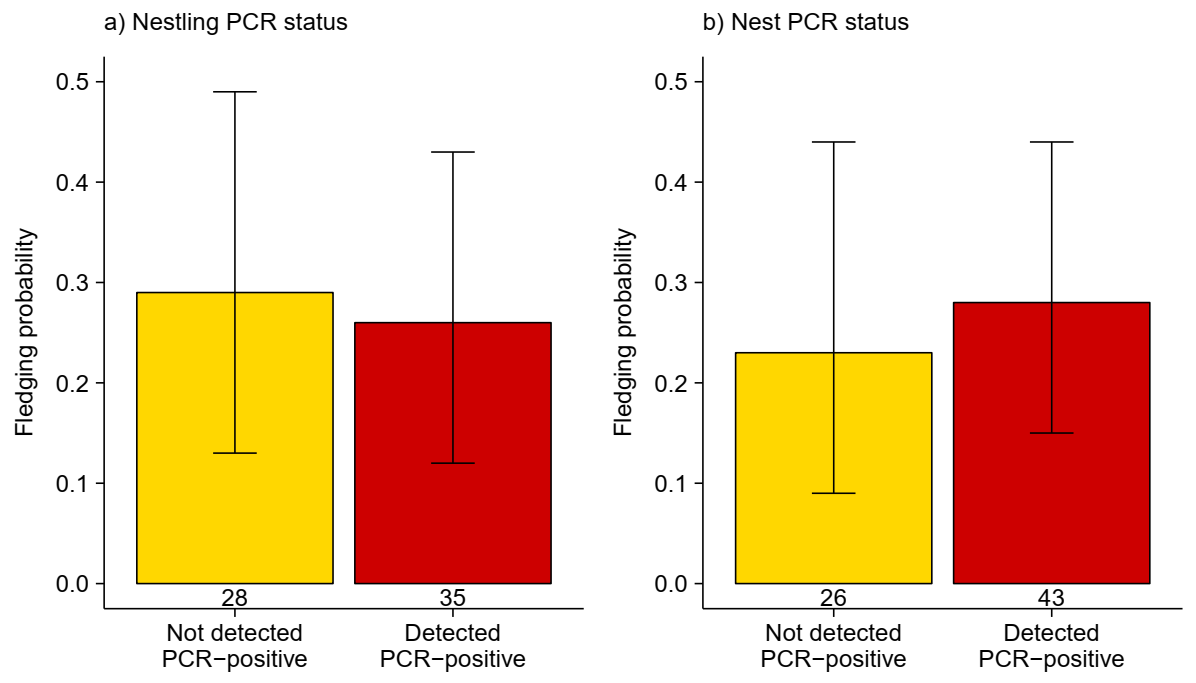

**Figure S5.** Relationship between fledging probability and *Pm­*-PCR status at the individual nestling (a) or whole nest (*i.e.*, including the parents’ samples; b) levels. The proportion of fledged nestlings are reported with their 95% Clopper-Pearson confidence intervals; sample sizes are indicated below the bars. A nestling was considered as “Detected PCR-positive” if *Pm* DNA was detected in at least one of its samples (a). At the nest level, a nest was considered as “Detected PCR-positive” if *Pm* DNA was detected in at least one of the nestling’s or parents’ samples (b). We could not detect any significant difference of the fledging probability between nestlings not detected as PCR-positive (proportion of fledged nestlings [95% confidence interval]: 0.29 [0.13; 0.49]) and nestlings detected PCR-positive (0.26 [0.12; 0.43]). The same result was obtained at the nest level (0.23 [0.09; 0.44] and 0.28 [0.15; 0.44] respectively).

**Interpretation and discussion.** Despite previous reports of a negative effect of *Pm* on albatross nestling survival on Amsterdam island (Weimerskirch, 2004; Bourret et al., 2018; Jaeger et al., 2018), we could not detect a correlation between *Pm*-PCR status and fledging probability at the individual or nest level in the context of this study. This seemingly contradictory observation is likely a consequence of a discrepancy between the temporal scales of the sampling (once a month) and that of the epidemiological process, avian cholera being considered as an acute disease with a high lethality rate in most cases (Wobeser, 1997). For instance, if a nestling is detected as PCR-negative during the February sampling occasion and dead at the following sampling, we cannot exclude the possibility that this nestling became infected between the two occasions. Estimating the lethality rate of *Pm* infection in the study population would require more frequent sampling occasions at the epizootic peak. The data collected as part of this study provide useful elements to optimize future field sampling designs.

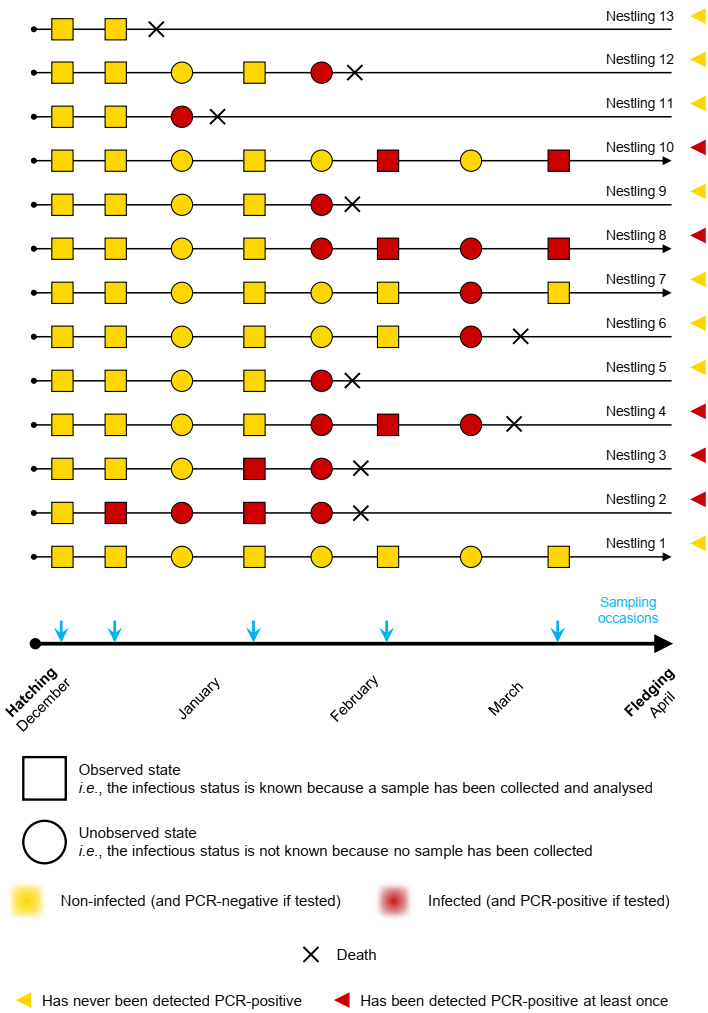

See next page for figure legend (Figure S6).

**Figure S6.** Schematic illustration of the importance of the timing of sampling for the inference of epidemiological processes. In the presented fictive situation, 11 nestlings got infected (dark red discs and squares; nestlings 2-12), among which 8 died following infection (nestlings 2-6, 9, 11 and 12), but only 5 were detected as PCR-positive (dark red triangles). Hence, more than half of the infected nestlings were “missed”. More precisely, in this fictive biological situation, 8/12 (67%) infected nestlings died *versus* 1/2 (50%) non-infected nestlings died; but the fictive study presented in this schema (following the sampling design illustrated with the blue arrows) would report that 3/5 (60%) PCR-positive nestlings died *versus* 6/8 (75%) PCR-negative nestling died leading to the false conclusion that infected nestlings have a lower mortality rate than non-infected nestlings. For illustrative purpose, we have chosen small sample sizes but the phenomenon would be the same with higher sample sizes as long as the timing of sampling relative to the epidemiological process is the same, potentially leading to statistical significant differences between infected and non-infected nestlings that do not represent the actual biological situation (which gives the opposite result). The same phenomenon (*i*.*e.*, discrepancy between the timing of the biological process and of sampling) will also interfere with the possibility to test other hypotheses such as the existence of a spatial structuration of the epidemiological dynamics (Figure S4).

This can occur if nestlings have time to get infected and die between two sampling occasions (e.g., in the case of acute diseases causing rapid deaths), they might never get detected as infected even if they actually got infected (e.g., nestlings 5, 9, 11 and 12). In the case of the present study, such issue could have arisen if the time delay between two sampling occasions (one month as of January) is longer than the delay between *Pm* infection and induced death, which is likely to be the case considering the pathogenicity of avian cholera (Wobeser, 2007; note however that the infectio period is not known). In addition, (1) non-infected nestlings can die from other causes (e.g., predation by skuas or rats, poor parental care…; e.g., nestling 13) and (2) infected nestlings can survive until fledging (as shown for the first time the present study; e.g., nestlings 7, 8 and 10). These two situations contribute to the blurring of the correlation signal between infection status and fledging probability. Hence the benefits of experimental approaches that allow to indirectly “control infection probability” (e.g., Bourret et al., 2018 in the same study system) and properly test causal links between two variables, which is not the objective of the present study.

Note that for the sake of simplicity, we did consider state misclassification (*i.e.*, sensitivity and specificity of test equal to 1), which is not likely to be the case, leading to additional blurring of the correlation signal between infection status and fledging probability. Designs accounting for uncertainty at all the stages of the observation process are called for (sampling and laboratory analyses; DiRenzo et al., 2018). The basic data we report in the present study will thus help to design future field studies in a iterative process (Restif et al., 2012), notably because such basic knowledge is essential to propose optimized repeated sampling designs that will help in precisely describing key epidemiological parameters, such as the probability to clean the infection, while accounting for the test detection probability (MacKenzie & Royle, 2005).

Overall, Figure S5, in the light of the explications given in this Figure S6, illustrates the importance of the timing of sampling in disease ecology studies and the necessity to describe epidemiological dynamics as a first step to design appropriate sampling designs to answer specific questions (see Yoccoz et al., 2001 for an analogy with biodiversity monitoring).

**Cited references**

Boulinier, T., Kada, S., Ponchon, A., Dupraz, M., Dietrich, M., Gamble, A., … McCoy, K. D. (2016). Migration, prospecting, dispersal? What host movement matters for infectious agent circulation? *Integrative and Comparative Biology, 56*, 330–342.

Bourret, V., Gamble, A., Tornos, J., Jaeger, A., Delord, K., Barbraud, C., … Boulinier, T. (2018). Vaccination protects endangered albatross chicks against avian cholera. *Conservation Letters, 11*, e12443.

BirdLife International (2018). *Diomedea amsterdamensis*. The IUCN Red List of Threatened Species 2018: e.T22698310A132397831.

Bredy, J. P., & Botzler, R. G. (1989). The effects of six environmental variables on *Pasteurella multocida* populations in water. *Journal of Wildlife Diseases, 25*, 232–239.

Caron, A., Cappelle, J., Cumming, G. S., de Garine-Wichatitsky, M., & Gaidet, N. (2015). Bridge hosts, a missing link for disease ecology in multi-host systems. *Veterinary Research, 46*, 83.

Curtis, P. E. (1983). Transmission of *Pasteurella multocida* infection from the brown rat (*Rattus norvegicus*) to domestic poultry. *Veterinary Record, 113*, 133–134.

DiRenzo, G. V., Campbell Grant, E. H., Longo, A. V., Che-Castaldo, C., Zamudio, K. R., & Lips, K. R. (2018). Imperfect pathogen detection from non-invasive skin swabs biases disease inference. *Methods in Ecology and Evolution, 9*, 380–389.

Gamble, A., Garnier, R., Jaeger, A., Gantelet, H., Thibault, E., Tortosa, P., … Boulinier, T. (2019). Exposure of breeding albatrosses to the agent of avian cholera: dynamics of antibody levels and ecological implications. *Oecologia*, in press.

Garnier, R., Ramos, R., Staszewski, V., Militão, T., Lobato, E., González-Solís, J., & Boulinier, T. (2012). Maternal antibody persistence: a neglected life-history trait with implications from albatross conservation to comparative immunology. *Proceedings of the Royal Society of London B: Biological Sciences, 279*, 2033–2041.

Haydon, D. T., Cleaveland, S., Taylor, L. H., & Laurenson, M. K. (2002). Identifying reservoirs of infection: a conceptual and practical challenge. *Emerging Infectious Diseases, 8*, 1468–1473.

Jaeger, A., Lebarbenchon, C., Bourret, V., Bastien, M., Lagadec, E., Thiebot, J.-B., … Weimerskirch, H. (2018). Avian cholera outbreaks threaten seabird species on Amsterdam Island. *PLOS ONE, 13*, e0197291.

Mackenzie, D. I., & Royle, J. A. (2005). Designing occupancy studies: general advice and allocating survey effort. *Journal of Applied Ecology, 42*, 1105–1114.

Pebesma, E.J. (2004). Multivariable geostatistics in S: the *gstat* package. *Computers & Geosciences, 30*, 683–691.

Samuel, M. D., Shadduck, D. J., Goldberg, D. R., & Johnson, W. P. (2003). Comparison of methods to detect *Pasteurella multocida* in carrier waterfowl. *Journal of Wildlife Diseases, 39*, 125–135.

Samuel, M. D., Shadduck, D. J., Goldberg, D. R., & Johnson, W. P. (2005). Avian cholera in waterfowl: the role of lesser snow and Ross’s geese as disease carriers in the Playa Lakes Region. *Journal of Wildlife Diseases, 41*, 48–57.

Weimerskirch, H. (2004). Diseases threaten Southern Ocean albatrosses. *Polar Biology, 27*, 374–379.

Wobeser, G. A. (1997). Avian cholera. In Diseases of wild waterfowl (2nd ed., pp. 57–69). Boston, MA: Springer US.

Yoccoz, N. G., Nichols, J. D., & Boulinier, T. (2001). Monitoring of biological diversity in space and time. *Trends in Ecology & Evolution, 16*, 446–453.
